# Protective Role of Podocytic IL-15 / STAT5 Pathway in Experimental Focal and Segmental Glomerulosclerosis

**DOI:** 10.1101/2022.11.07.515420

**Authors:** Aïssata Niasse, Kevin Louis, Olivia Lenoir, Chloé Schwarz, Xiaoli Xu, Aymeric Couturier, Hélène Dobosziewicz, Anthony Corchia, Sandrine Placier, Sophie Vandermeersch, Lothar Hennighausen, Perrine Frere, Pierre Galichon, Brigitte Surin, Souhila Ouchelouche, Liliane Louedec, Tiffany Migeon, Marie-Christine Verpont, David Buob, Yi-Chun Xu-Dubois, Hélène Francois, Eric Rondeau, Laurent Mesnard, Juliette Hadchouel, Yosu Luque

**Author notes:** **Corresponding author:** Dr. Yosu LUQUE, M.D., Ph.D., Soins Intensifs Néphrologiques et Rein Aigu. Département de Néphrologie. Sorbonne Université. Tenon Hospital, 4 rue de la Chine, Paris, France 75020. **Sources of support:** ANR JC (Agence Nationale de la Recherche, Jeune Chercheur) PODOGAMMAC - ANR-17-CE14-0004 French Society of Nephrology (SFNDT) grant for translational research project on podocytic diseases (2017).

## Abstract

During glomerular diseases, podocyte-specific pathways can modulate the intensity of the lesions and prognosis. The therapeutic targeting of these pathways could thus improve the management and prognosis of chronic kidney diseases. The Janus Kinase/ Signal Transducer and Activator of Transcription (JAK/STAT) pathway, classically described in immune cells, has been recently described in intrinsic kidney cells. Here, we show, for the first time, that STAT5 is activated in human podocytes in focal segmental glomerulosclerosis (FSGS). Additionally, *Stat5* podocyte-specific inactivation aggravates the functional and structural alterations in a mouse model of FSGS. This could be due, at least in part, to an inhibition of the autophagic flux. Finally, Interleukin 15 (IL-15), a classical activator of STAT5 in immune cells, increases STAT5 phosphorylation in human podocytes and its administration alleviates glomerular injury *in vivo* by maintaining the autophagy flux in podocytes. In conclusion, activating podocytic STAT5 with commercially available IL-15 represents a new therapeutic avenue with the potential for FSGS.

## Introduction

Podocytes, specialized epithelial cells of the glomerulus, are one of the main targets of injury in kidney diseases. A range of injuries (immune, toxic, hypertensive) affects podocytes, thus driving the progression to chronic kidney disease (CKD) and end-stage kidney disease (ESKD), both of which are increasing in global frequency. Current specific therapies for CKD are limited and ESKD treatment is based mostly on replacement therapies such as dialysis or transplantation that are non-optimal and associated with several side effects and a decrease in the quality of health and life. The importance of tissue-specific effectors, acting in kidney epithelial cells, vessels, or the renal interstitium, has been highlighted by several studies over the past few years^1–3^. Their modulation can significantly modify the course and evolution of kidney injury towards CKD and ESKD. Thus, the therapeutic targeting of these pathways could improve CKD management and prognosis.

Classically described in immune cells, the Janus Kinase-Signal Transducer and Activator of Transcription (JAK-STAT) pathway, has been shown to be important in the kidney in recent years^4^. The JAK/STAT system regulates cell differentiation, growth and survival. It plays an important role in immune cells, where it is activated by several cytokines. However, it is also expressed in non-immune cells^5^. For example, STAT3 in podocytes contributes to glomerular injury in experimental glomerulonephritis^6,7^ and to podocyte proliferation in experimental HIV-associated nephropathy^8^. Additionally, STAT3 activation in tubular cells leads to lipocalin-2 and PDGFβ production with subsequent matrix expansion and fibrosis^9^.

We have previously shown that the common gamma chain (γC, encoded by the *Il2rg* gene), an interleukin subunit receptor that activates the JAK-STAT pathway, is expressed *de novo* by podocytes during human crescentic glomerulonephritis^1^. γC kidney-deficiency aggravates experimental glomerulonephritis in a lymphocyte-independent manner, suggesting a protective role of the γC/JAK/STAT system in kidney cells. Moreover, interleukin-15 (IL-15) is able to activate downstream JAK/STAT signaling in cultured podocytes in a γC-dependent manner. STAT5 is a transcription factor with two isoforms (STAT5A and STAT5B), activated by γC/JAK signaling and expressed by epithelial cells^10^. STAT5 is transactivated by tyrosine phosphorylation and classically regulates proliferation and apoptosis pathways^11,12^. In mice, complete *Stat5* inactivation causes perinatal lethality and reduced development of B- and T-cells^13^. Given the protective role of renal γC during experimental glomerulonephritis, we hypothesized that activation of podocytic STAT5 would be protective in glomerular diseases.

We observed an upregulation of glomerular STAT5 in FSGS. Deficiency of STAT5 in podocytes aggravated kidney injury in inflammatory and non-inflammatory glomerular disease mouse models. This could be due, at least in part, by an inhibition of the autophagic flux in podocytes. Therapeutic stimulation of STAT5 *in vivo* by IL-15 alleviated experimental FSGS and maintained podocytic autophagic flux. Our findings provide a potentially novel target for glomerular disease.

## Materials and Methods

### Human tissues

Paraffin embedded renal biopsy specimens were obtained from the Tenon hospital [Assistance Publique - Hôpitaux de Paris (AP-HP), Paris, France]. Human tissue was used after informed consent by all patients and signature of an approval form. The kidney biopsy collection was approved by the Ethics Committee of the AP-HP. Kidney biopsy specimens with sufficient tissue for immunohistochemical evaluation after the completion of diagnosis workup were included. We analyzed kidney biopsies from patients with the following diagnosis: HIV associated nephropathy (HIVAN) (n=3), non-HIVAN collapsing FSGS (n=4) and non-collapsing FSGS (n=3). 3-months post-transplant kidney protocol biopsies without any histological abnormality according to 2017 Banff classification were used as controls (n=3).

### Animals

All procedures regarding animal experimentation were conducted in accordance with the European Union Guidelines for the Care and Use of Laboratory Animals and approved by the local ethics committee of the National Institute for Health and Medical Research (Institut National de la Santé et de la Recherche Médicale, Inserm). Animals were housed at a constant temperature with free access to water and food.

Mice with a podocyte-specific disruption of *Stat5a* and *Stat5b* genes (*Nphs2*.*cre*-*Stat5^lox/lox^*) were generated by crossing *Nphs2*.*Cre* mice^14^ with *Stat5^lox/lox^* (kindly provided by L.Hennighausen^15^) on the C57BL6/J background. *Nphs2*.cre-*Stat5*^lox/lox^ mice and control littermate, aged 10 to 12 weeks, were used in this study.

BALB/C mice were obtained from Janvier Labs (Le Genest-Saint-Isle, France). 10-12 weeks-old mice were used for the experiments.

### Experimental adriamycin nephropathy

Adriamycin (doxorubicin) (Adriblastine 50 mg, Pfizer, France) or saline (controls) was injected intravenously in 10-12 weeks-old mice at day 0. As previously described^16^, we used a 25 mg/kg dose for experiments with C57BL/6J mice (as that genetic background is naturally resistant to adriamycin) and 10 mg/kg for experiments with BALB/C mice. To prevent C57BL/6J mice from excessive weight loss and malnutrition, 1mL of glucose peritoneal dialysis fluid (Physioneal, Baxter Healthcare Ltd, US) was intraperitoneally injected daily to the mice, as previously described^16^. The mice were weighed at day 0, 2, 3, 4, 6 and 7. Urine samples were collected at day 0 and day 7. At day 7, mice were anesthetized with a mix of ketamine and xylazine (10 and 0.1mg/kg) and blood samples were collected from the retro-orbital sinus. Mice were then euthanized and kidneys and spleen were collected for analysis.

### Experimental anti-GBM glomerulonephritis

Passive experimental anti-GBM glomerulonephritis was induced using decomplemented sheep anti-rat GBM serum prepared as previously reported^17^. Passive non-accelerated anti-GBM glomerulonephritis was induced by intravenous administration of 1.5 mg total protein/g body weight over three consecutive days. Renal injury was evaluated on day 9 following the first injection of anti-GBM serum. As controls, mice were injected with phosphate buffered saline.

### Renal function and proteinuria assessment

Plasma urea and urine protein and creatinine levels were measured with a Konelab enzymatic spectrophotometric analyzer (ThermoFischer Scientific, Waltham, MA, US). Urine albumin-to-creatinine ratio were measured using a Beckman Coulter AU 480 analyzer.

### Hemograms

Blood samples were analyzed with IDDEX ProCyte Dx™ hematology analyzer kindly provided by Dr Ivan Moura and Michael Dussiot (Inserm U1163 / CNRS ERL 8254, Institut Imagine, Paris, France).

### Quantitative RT-PCR

RNA was extracted from whole kidneys using TRIzol solution (Life Technologies). Residual genomic DNA was removed by DNase I treatment (Thermo Scientific™). 1μg of total RNA was reverse transcribed using the Maxima First Strand cDNA Synthesis Kit for RT-qPCR (Thermo Scientific™). HPRT was used as endogenous reference housekeeping gene. The specific PCR primer sequences used are listed in **Supplemental Table 1**. The qPCR reaction was performed using a LightCycler 480 (Roche Diagnostics). The analysis was performed using the 2-ΔΔC_t_ method. Results are expressed as arbitrary units, which represent the ratio of the target gene to the internal control gene (HPRT).

### Histological and immunohistochemical studies

Kidneys were fixed in formalin solution (4%) and embedded in paraffin. Sections (3 μm thick) were processed for histopathology study or immunohistochemistry. Sections were stained with Masson’s trichrome, Periodic acid–Schiff (PAS) or Martius Scarlet Blue (Microm Microtech®).

Tubular lesions (tubular dilation, loss of brush border, cast formation) were evaluated semi quantitatively using Masson’s trichrome or PAS stained sections as follows: 0, no lesion; 1, lesions in 1% to 25% of the tubules analyzed; 2, lesions in 26% to 50% of the tubules analyzed; 3, lesions in 51% to 75% of the tubules analyzed; 4, lesions in >76% the tubules analyzed. Tubular injury score was evaluated in 10 randomly selected, non-overlapping fields at objective 20x in each section by two blinded observers.

For immunohistochemistry, paraffin-embedded sections were stained with the following primary antibodies: rabbit anti-phospho-STAT5 B (Abcam, ab52211, 1:100), rabbit anti-CD3 (Abcam, ab5690, 1:150), rat anti-F4/80 (Biorad, MCA97R, 1:200), recombinant anti-Cre (Merck Millipore, 69050, 1:1000) and revealed using Histofine reagent (Nichirei Biosciences). Human kidney biopsies were stained with rabbit anti-phospho-STAT5 B (Abcam) and revealed using Histofine reagent (Nichirei Biosciences).

For nephrin immunofluorescence, 3μm-thick frozen sections were stained with the following antibodies: anti-nephrin (AF3159, R & D Systems, 1:200). The signal was detected with Alexa Fluor secondary antibodies followed by DAPI counterstaining and mounting in PermaFluor medium (Thermo Fisher Scientific). For P62 paraffin embedded tissue immunofluorescence we used anti-SQSTM1/P62 (1:1000, PROGEN, GP-62C), goat anti-Podocalyxin (PODXL; 1:1000, Bio-Techne, AF-1556). Secondary antibodies were Alexa 488– and Alexa 568–conjugated antibodies from Invitrogen. Nuclei were stained in blue using Hoechst. Slides were mounted using fluorescent mounting medium (Dako, S3023). Photomicrographs were taken with a Zeiss Axiophot photomicroscope and Axiovision software. Semiautomatic quantifications on Fiji were used for quantifications of PODXL+ areas per glomerular section on at least 30 glomeruli per mouse.

The proportion of glomeruli with fibrin deposits was evaluated by examination of at least 30 glomeruli per cortical section for each mouse after Martius Scarlet Blue staining, by an examiner (A.N.) who was blinded to the experimental conditions.

ImageJ software (National Institutes of Health) was used for the measurement of the surface of glomerular nephrin-positive staining over total glomerular area, number of CD3 or F4/80 positive cells over section area and KIM-1 positive area over section area. Widefield and confocal microscopy were performed using an inverted microscope IX83 (Olympus, Tokyo, Japan) equipped with DSD2 system attached to an Andor Zyla camera (Andor Technology, Belfast, UK).

### Electron Microscopy

Kidneys were cut into small pieces, and immersed in 2.5% glutaraldehyde containing 1% tannic acid in 0.1 mol/L PBS for 2 hours at 4°C. Samples were post-fixed with 1% OsO4, dehydrated, and embedded in epoxy resin. Ultrathin sections were stained with uranyl acetate and lead citrate and then examined under a Philips CM10 electron microscope (Philips Innovation Services, Eindhoven, the Netherlands).

### Isolation of glomeruli and primary cultured podocytes

Freshly isolated renal cortexes from *Stat5^lox/lox^* and *Nphs2.Cre-Stat5^lox/lox^* mice were mixed and digested by collagenase I (1 mg/ml; Invitrogen) in RPMI 1640 (Invitrogen) for 2 minutes at 37°C. Next, collagenase I was inactivated with RPMI 1640 plus 10% FBS (Hyclone). Tissues were then passed through a 100-μm cell strainer and a 40-μm cell strainer (BD Falcon) in PBS (Euromedex) with 0.5% BSA (Euromedex). Glomeruli, adherent to the 40-μm cell strainer, were removed with PBS with 0.5% BSA injected under pressure, and finally washed twice in PBS. Freshly isolated glomeruli purity was analyzed by light microscopy and then were plated in six-well dishes in RPMI 1640 with 10% FBS, 2% HEPES buffer (Gibco), and 1% penicillin/streptomycin (Gibco). Two days after seeding, glomeruli became adherent to the dish, and podocytes spread out from glomeruli. Primary podocytes were then mixed in RIPA buffer (Santa Cruz) with orthovanadate, PMSF, protease inhibitor cocktail (Santa Cruz) and NaF and frozen at −80°C for protein extraction.

*Western blot analysis (see Supplementary Methods)*

*Cell Culture (see Supplementary Methods)*

*Generation of STAT5B knock-out in immortalized human podocytes (see Supplementary Methods)*

*IL-15 therapy protocol in vivo (see Supplementary Methods)*

### Statistics

All data are expressed as means ± SEM. Statistical analysis was performed using the GraphPad Prism software. The Mann-Whitney test was used to compare two groups. The Kruskal-Wallis test was used to analyze the distribution of three or more groups. A value <0.05 was considered significant.

## Results

### STAT5 activation in podocytes in human focal & segmental glomerulosclerosis (FSGS)

We explored the pathways activated by γC in human glomerular kidney diseases (**Supplemental Table 2**). STAT5A and STAT5B are activated by phosphorylation and nuclear translocation. Therefore, we performed an immunostaining against phosphorylated STAT5A and STAT5B (pSTAT5) on kidney biopsies from patients with both collapsing and non-collapsing FSGS. The former is a severe form characterized by apoptosis and proliferation of injured podocytes. We found no differences in pSTAT5A between controls and disease biopsies (data not shown). In contrast, there was a significant accumulation of pSTAT5B in the nucleus of podocytes of patients (**Figure 1**). Interestingly, STAT5B activation was more intense in collapsing forms of FSGS than in non-collapsing forms. Additionally, nuclei expressing higher amounts of pSTAT5B appeared enlarged.

**Figure 1.**
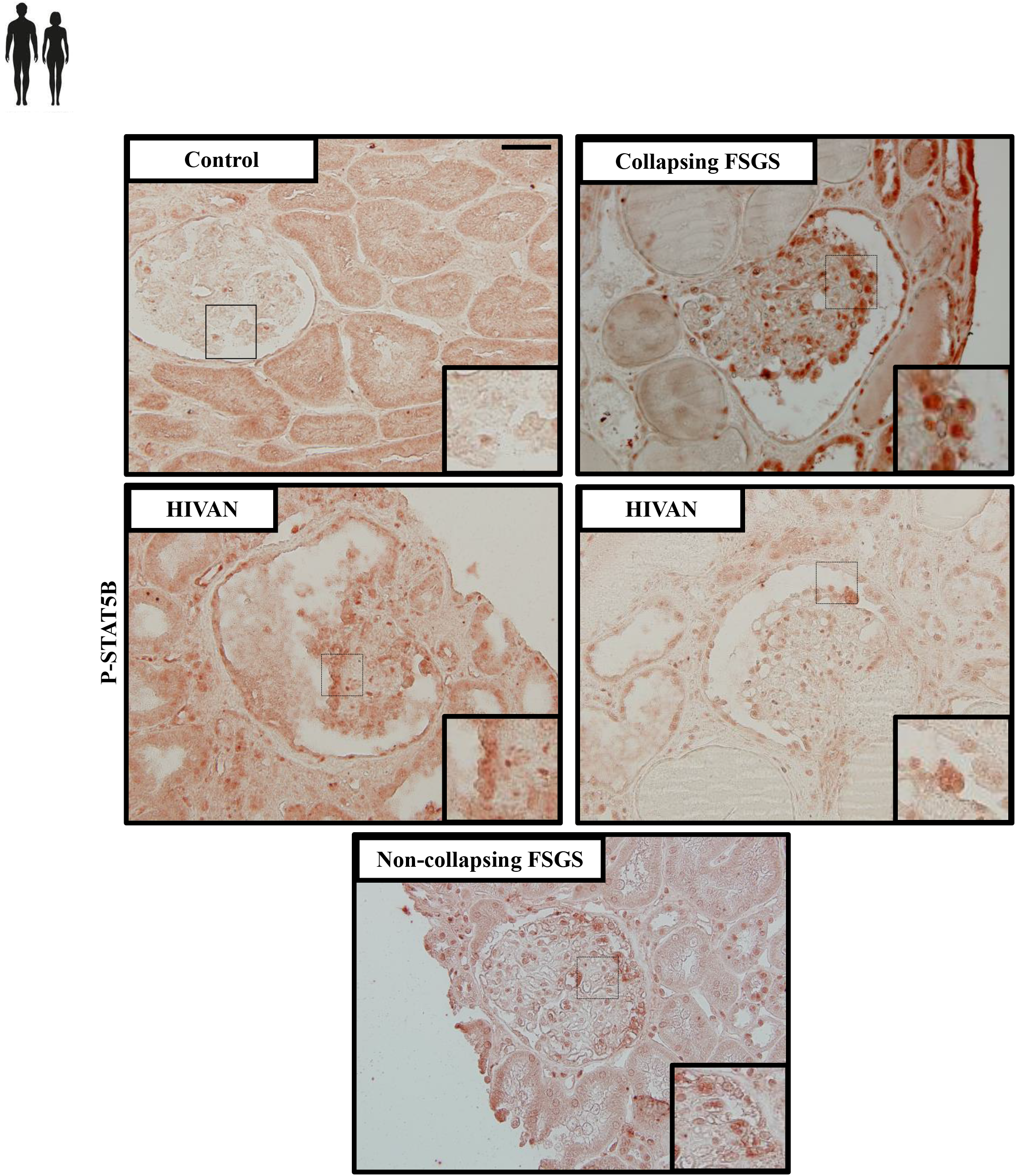
STAT5 activation in podocytes in human focal & segmental glomerulosclerosis (FSGS) Representative images of phospho-STAT5B immunostaining in human kidney biopsies from patients with HIVAN, non-HIVAN collapsing, non-collapsing FSGS and pristine 3 months post-transplant kidney biopsy as a control. Scale bar, 50μm. Higher magnifications are shown in the insets. Each microphotograph corresponds to an individual patient’s biopsy.

### STAT5 deficiency in podocytes does not affect renal development in mice

To further examine the role of STAT5 in podocytes, we generated a podocyte-specific knockout of *Stat5* (*Nphs2-Cre;Stat5^lox/lox^*) (**Supplemental Figure 1a**). We confirmed podocyte-specific STAT5 deficiency by immunoblotting of primary mouse podocytes isolated from control and *Nphs2-Cre;Stat5^lox/lox^* kidneys (**Figure 2a**) and podocyte-specific Cre recombinase expression in *Nphs2.Cre-Stat5^lox/lox^* mice compared to control *Stat5^lox/lox^* mice using immunohistochemistry (**Supplemental Figure 1b**). *Nphs2-Cre;Stat5^lox/lox^* mice were viable and fertile. The renal function and structure were similar to those of *Stat5^lox/lox^* littermates (**Supplemental Figures 1b-d**), suggesting that STAT5 is not essential for podocytes development and function.

**Figure 2.**
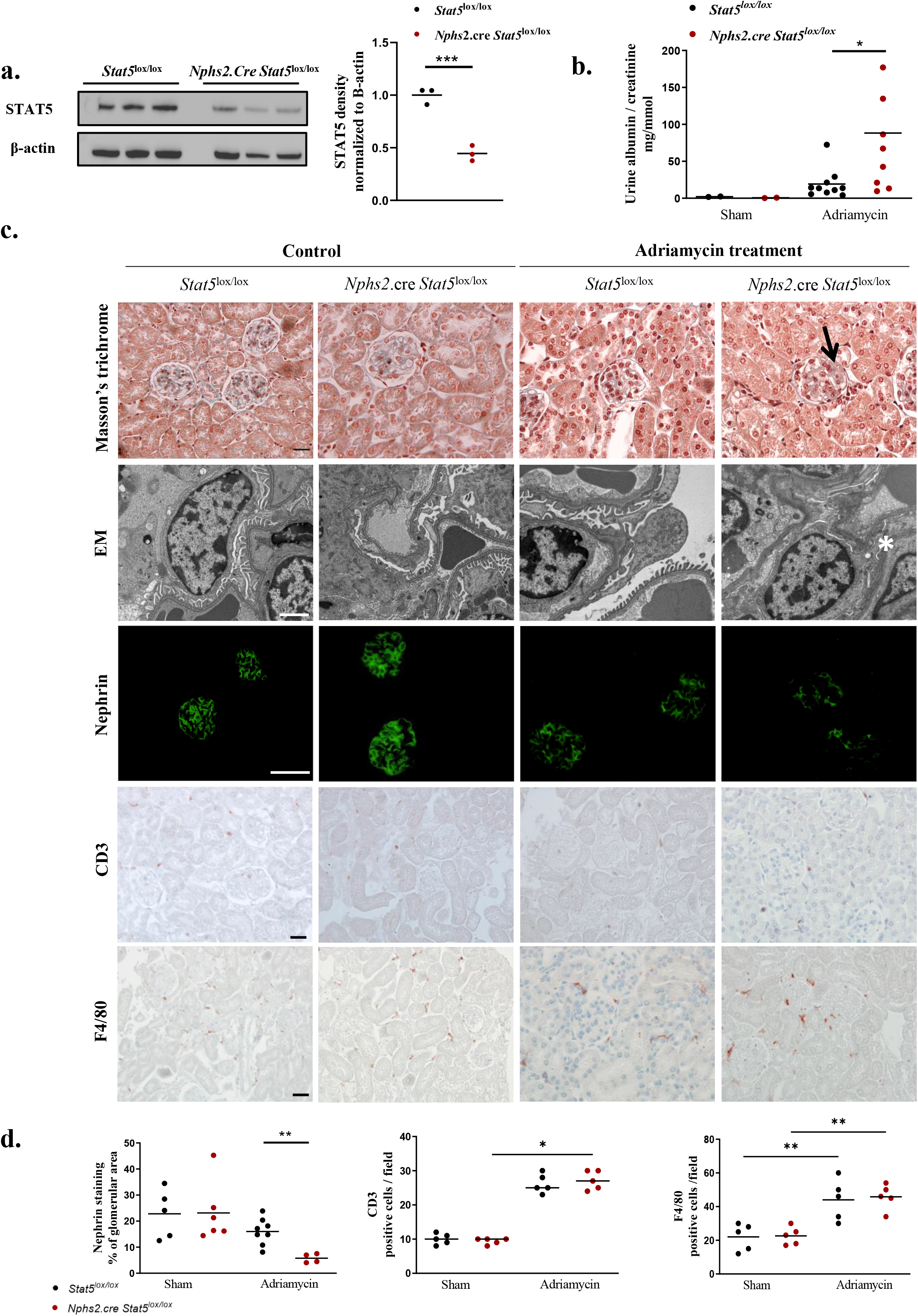
Podocyte STAT5 deficiency aggravates adriamycin-induced albuminuria and glomerular injury. **(a)** Western blot analysis of STAT5 expression in primary cultured podocytes from glomeruli isolated from 10-week-old *Stat5^lox/lox^* and *Nphs2*.cre-*Stat5^lox/lox^* mice. Beta-actin was used as loading control. Quantification of western blot bands for STAT5 normalized to Beta-actin band intensity. ***p<0.001 **(b)** Urinary albumin-to-creatinine ratio in *Stat5^lox/lox^* and *Nphs2*.cre-*Stat5^lox/lox^* mice 7 days after adriamycin or vehicule injection (sham group). Individual values are shown and the bars correspond to means. *p < 0.05. **(c)** Representative images of Masson’s trichrome stained sections (upper panels, scale bar: 20μm), transmission electron micrographs (second row panels, scale bar 2μm), nephrin staining (third row panels, scale bar, 20μM), CD3 staining (fourth row panels, scale bar 20μm) and F4/80 staining (fifth row panels, scale bar 20μm) of the renal cortex from *Stat5^lox/lox^* and *Nphs2*.cre-*Stat5^lox/lox^* mice 7 days after the injection of adriamycin. Arrow indicates glomerulosclerosis observed in optical microscopy and * indicates podocyte feet effacement and glomerular basement membrane enlargement observed with EM in *Nphs2*.cre-*Stat5^lox/lox^* mice. **(d)** Quantification of nephrin immunostaining, CD3 and F4/80 positive cells in kidneys from *Nphs2*.cre-*Stat5^lox/lox^* mice compared to *Stat5^lox/lox^* mice at day 7 after adriamycin injection. Individual values are shown and the bars correspond to means. *p < 0.05. **p<0.01.

### Podocyte STAT5 deficiency aggravates adriamycin-induced albuminuria and glomerular injury

To determine the role of podocyte STAT5 in glomerular injury, we induced adriamycin (doxorubicin) nephropathy, a model of FSGS, in *Nphs2-Cre;Stat5^lox/lox^* and *Stat5^lox/lox^* mice. In both groups, adriamycin induced a similar and significant weight loss (**Supplemental Figure 2a)** with a dramatic decrease in circulating leukocytes and platelets, but not hemoglobin (**Supplemental Figure 2b**). 7 days after Adriamycin administration, podocyte STAT5 deficiency was associated with a 5-fold increase in proteinuria (**Figure 2b**) and more severe glomerular lesions (**Figure 2c**). Electron microscopy (EM) demonstrated podocyte flattening, foot process effacement and glomerular basement membrane thickening in *Nphs2-Cre;Stat5^lox/lox^* mice **(Figure 2c)**. Additionally, the expression of nephrin was reduced in STAT5-deficient mice compared to controls (**Figures 2c and 2d**). Adriamycin increased renal T-cell and macrophage infiltration but these were unaffected by podocyte STAT5 deficiency (**Figures 2c and 2d**). Taken together, these results suggest that podocyte STAT5 protects the glomeruli from injury.

### Podocyte STAT5 deficiency aggravates proteinuria and glomerular lesions in anti-glomerular basement membrane (GBM) experimental glomerulonephritis

Next, we induced anti-GBM glomerulonephritis in podocyte-specific Stat5-deficient mice. Whereas podocyte Stat5 deficiency was associated with increased proteinuria (**Figure 3a**), glomerular injury and ultrastructural abnormalities (**Figure 3b and 3c**), the degrees of acute kidney injury and tubulo-interstitial disease were similar between control and mutant mice (**Figure 3c to 3e**). Taken together, these two models demonstrate that podocyte STAT5 activation protects against glomerular injury.

**Figure 3.**
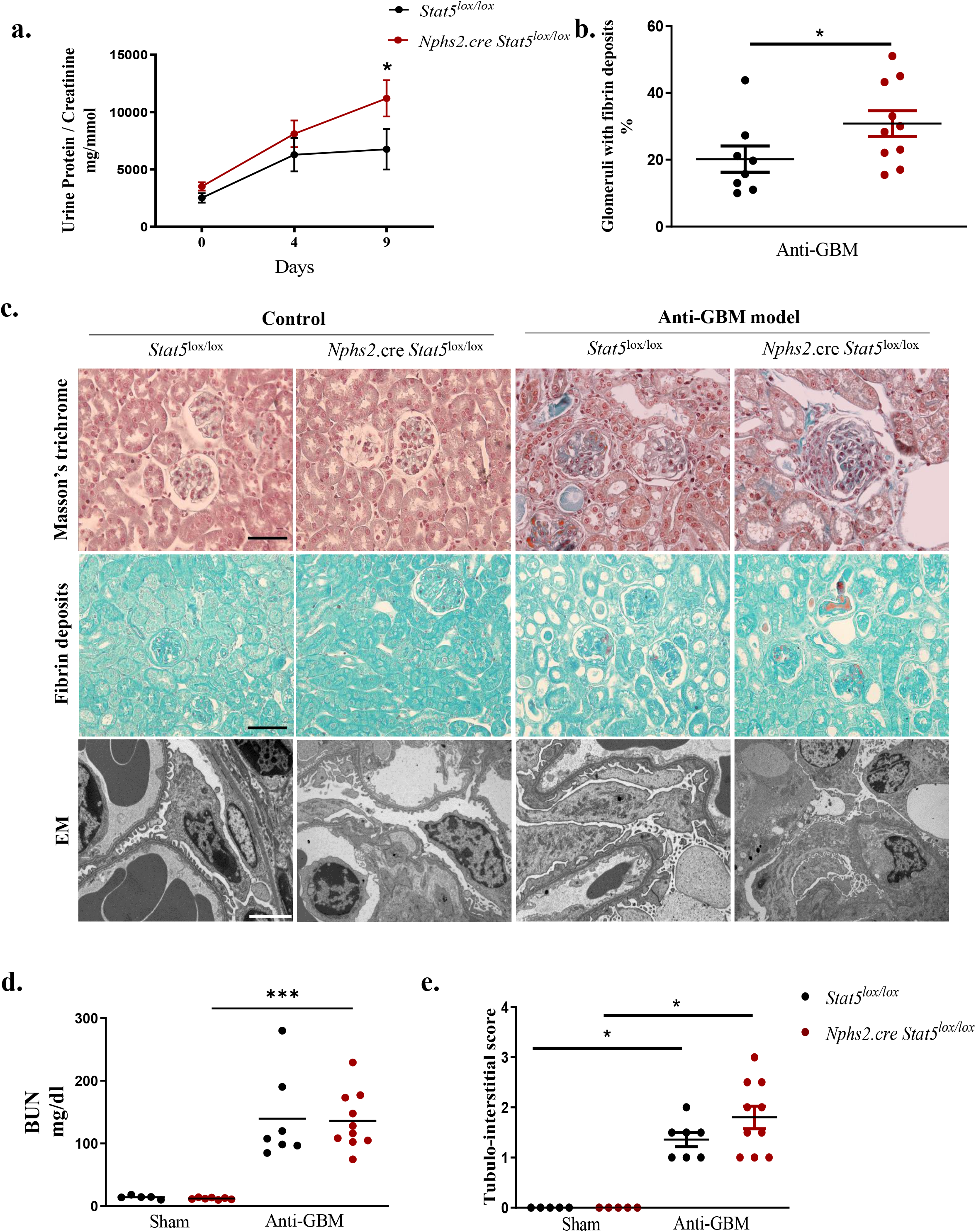
Podocyte STAT5 deficiency aggravates proteinuria and glomerular lesions in anti-glomerular basement membrane (GBM) experimental glomerulonephritis. **(a)** Urine protein-to-creatinine ratio at day 0, 4 and 9 after anti-GBM serum injection in *Stat5^lox/lox^* and *Nphs2*.cre*-Stat5^lox/lox^* (n= 7 to 9 mice per group). Data are mean ± s.e.m. *p <0.05. **(b)** Quantification of fibrin glomerular deposits in kidneys from *Stat5*^lox/lox^ and *Nphs2*.cre*-Stat5^lox/lox^* mice at day 9 after anti-GBM serum injection. Individual values are shown and the bars correspond to the means. *p<0.05. **(c)** Representative images of Masson’s trichrome-(upper panel), Martius Scarlett Blue-stained sections showing glomerular fibrin deposits (middle panel) and transmission electron micrographs (lower panel) of renal cortex from 10–12-week-old anti-GBM-glomerulonephritis-induced *Stat5*^lox/lox^ and *Nphs2*.cre*-Stat5^lox/lox^* mice and controls. Scale bar, 20μM for upper panel, 20μm for middle panel and 2μm for lower panel **(d)** Plasma BUN in *Stat5^lox/lox^* and *Nphs2*.cre*-Stat5^lox/lox^* mice after vehicle or anti-GBM injection. Individual values are shown and the bars correspond to the means. ***p <0.001, **(e)** Quantification of tubulo-interstitial damage in kidneys from *Stat5*^lox/lox^ and *Nphs2*.cre*-Stat5^lox/lox^* mice at day 9 after anti-GBM serum injection. Individual values are shown and the bars correspond to the means. *p<0.05.

### Interleukin-15 administration alleviates experimental FSGS in mice

IL-15 activates STAT5 through γC signaling in immune cells^18^. However, IL-15 is also produced by the renal epithelium^19^ and we have previously shown that IL-15 activates JAK signaling in podocytes^1^. Therefore, we tested if IL-15 could activate STAT5 in human podocytes *in vitro*. Thirty minutes of exogenous IL-15 stimulation increased STAT5B phosphorylation in cultured human podocytes (**Figure 5a**).

We went on studying the therapeutic effects of IL-15 administration in glomerular disease. The covalent binding of IL-15 to soluble IL-15Rα mimics trans-presentation and increases the bio-stability of IL-15 and its efficacy *in vivo*^20^. In adriamycin-induced FSGS, the administration of the IL-15/IL-15Rα complex (24h prior to, and at 1, 3 and 5 days following adriamycin) increased renal STAT5A and STAT5B mRNA expression (**Figure 5b**), with a trend towards increased STAT5 phosphorylation (**Figure 5c**). In accordance with the hypothesis that STAT5 activation could be protective, we observed that IL-15/IL-15Rα administration strongly reduced adriamycin-induced albuminuria and podocyte ultrastructural abnormalities in mice (**Figures 5d and 5e**).

IL-15 administration alone was not associated with any renal effects, although we did observe splenomegaly, in keeping with a role for IL-15 in leucocyte proliferation (**Supplemental Figure 3a**). Whereas IL-15/IL-15Rα administration did not modify MCP1 mRNA expression and renal macrophage infiltration, the number of T-cell present in the kidney increased (**Supplemental Figure 3b, c and d**).

### Modulation of the autophagic flux by STAT5 and IL-15 in vitro and in vivo

Autophagy has been shown to be protective in mouse models of glomerular diseases and acute kidney injury (AKI)^21–23^. Importantly, it is downregulated in several human glomerular pathological conditions^22–24^. Autophagy is a critical homeostatic pathway of podocytes, characterized by the removal of dysfunctional organelles and misfolded proteins by enclosure within double membrane vesicles, called autophagosomes, and their subsequent lysosomal fusion and degradation.

In order to characterize the potential regulation of the podocyte autophagic flux by STAT5, we generated human immortalized podocytes with a genetic deletion of *STAT5B* using the Crispr-Cas9 system (**Figure 4a**). The autophagic flux was measured by quantifying the level of expression of LC3b, following serum deprivation and exposure to bafilomycin. As shown in **Figure 4b**, the autophagic flux in *STAT5^−/-^* podocytes was decreased compared to *STAT5^+/+^* ones, suggesting that the inhibition of autophagy could be, at least in part, responsible for the severe adriamycin-induced renal alterations observed in mice when STAT5 is inactivated in podocytes.

**Figure 4:**
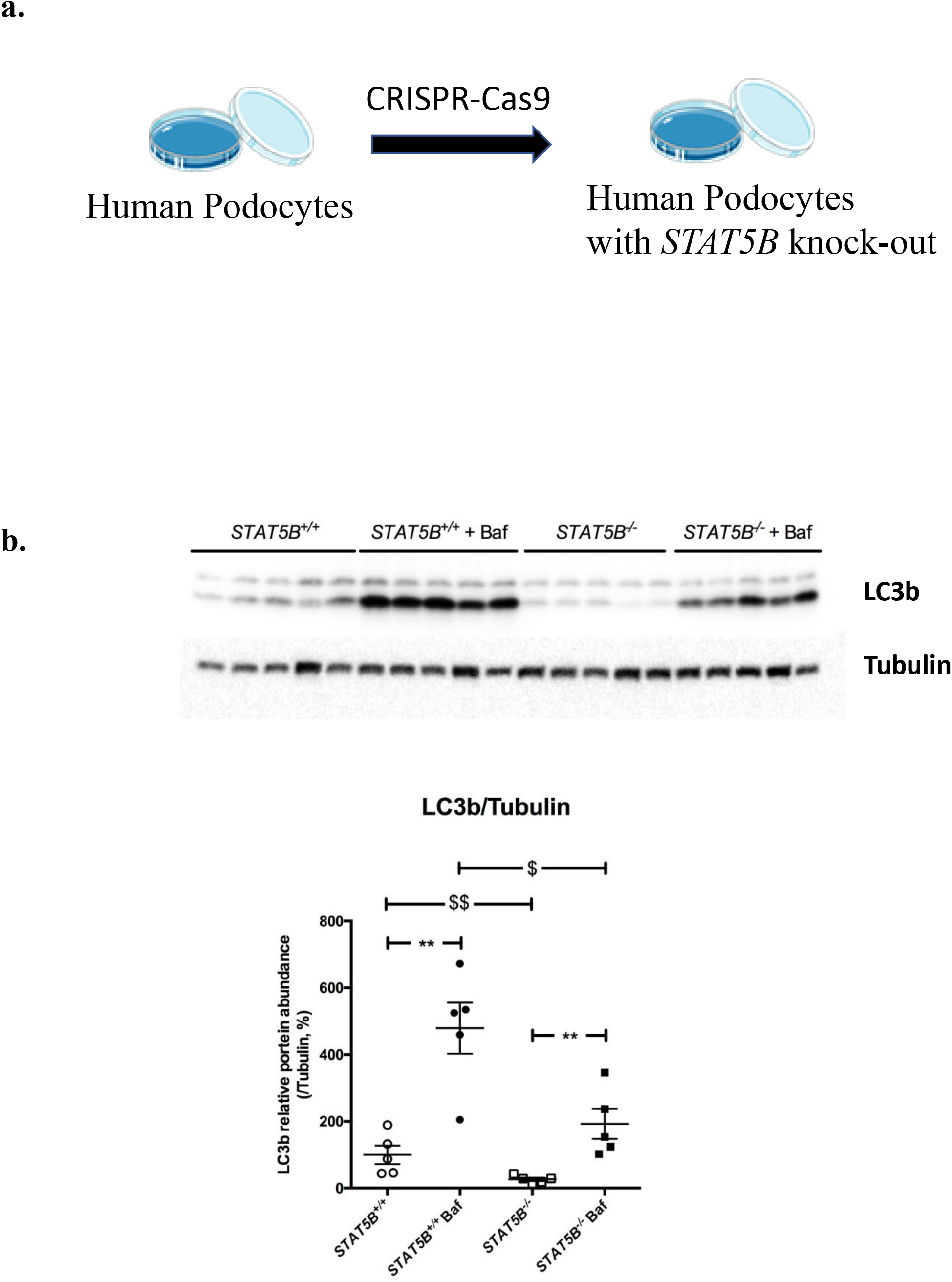
Blocking autophagosomal degradation confirmed decreased autophagic flux in STAT5B deficient human podocytes. **(a)** Scheme of CRISPR-Cas9 mediated *STAT5B* knock out in human podocytes. **(b)** Western blot analysis of the expression of LC3b in human podocytes with or without STAT5B deficiency. Tubulin expression serves as normalization. Podocytes were treated or not treated with bafilomycin A1 (BafA1; 100 nM) for 4 hours before culture arrest.

We then evaluated the level of autophagy in mice glomeruli using P62 immunostaining *in vivo* and as expected, we observed an increase P62 staining, reflecting a decrease autophagy, in the glomeruli of adriamycin-exposed mice. Interestingly, this staining decreased in the glomeruli of IL-15 treated mice (**Figure 5f and g**), suggesting that IL-15 administration improves the glomerular autophagic flux.

**Figure 5.**
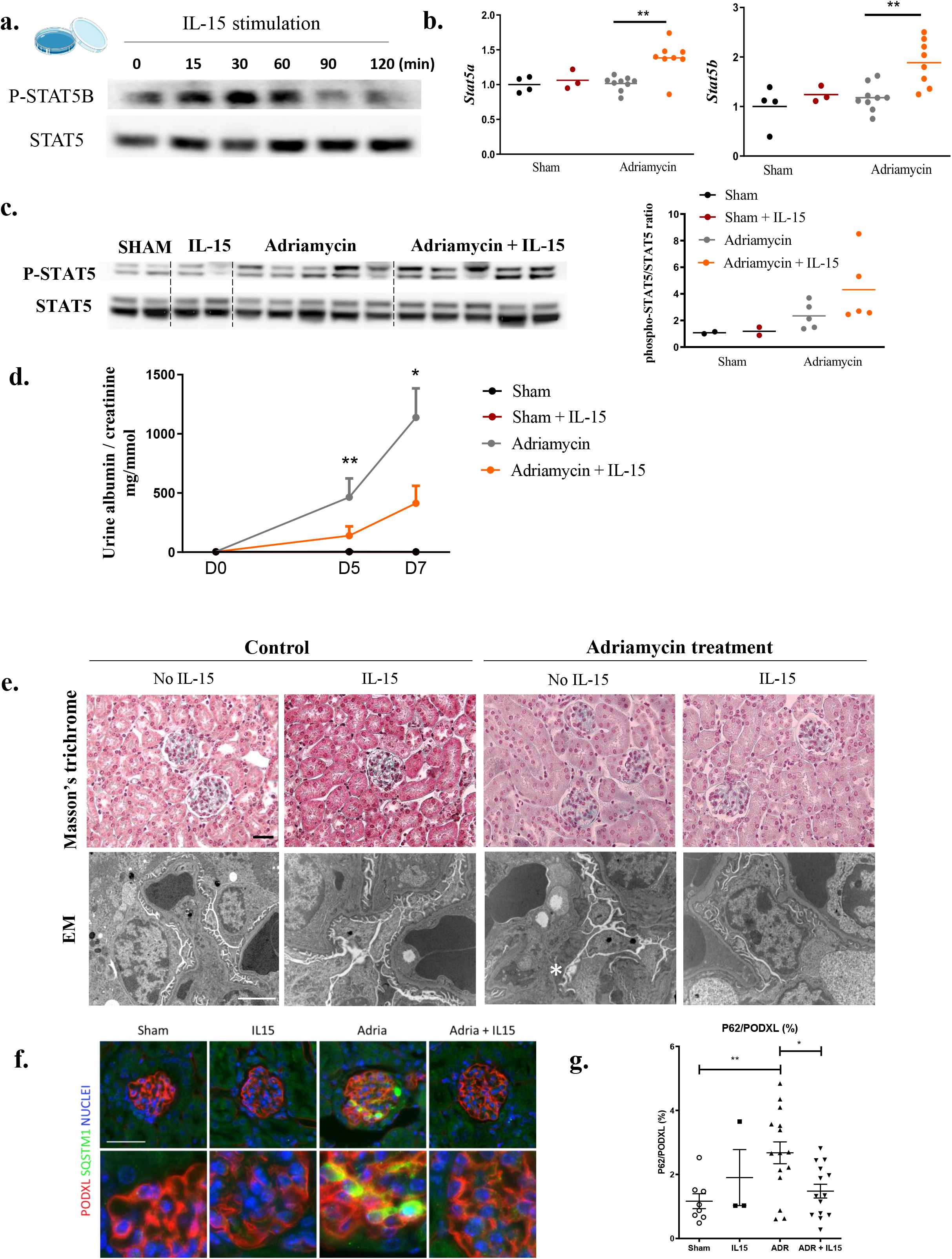
Interleukin-15 activates STAT5 in podocytes, modulates autophagy and alleviates glomerular injury. **(a)** Western blot analysis of the expression of phospho-STAT5B and STAT5 in human podocyte cell line after IL-15 stimulation at different time points. **(b)** RT-qPCR quantification of the expression of *Stat5a* and *Stat5b* in the kidneys from 12-week-old BALB/C mice at day 7 after adriamycin injection with or without IL-15 treatment. Individual values are shown and the bars correspond to the means. **p<0.01. **(c)** Western blot analysis of the expression of phospho-STAT5 and STAT5 in the kidneys from 12-week-old BALB/C mice at day 7 after adriamycin injection with or without IL-15 treatment and its quantification. Individual values are shown and the bars correspond to the means. **(d)** Urine albumin-to-creatinine ratio at day 0, day 5 and day 7 after adriamycin injection in BALB/C mice treated or not with IL-15. The data represent means ± s.e.m. *p< 0.05. **p< 0.01. **(e)** Representative images of Masson trichrome (upper panel; scale bar: 20μM) and transmission electron micrographs (lower panel; scale bar : 2μm) of kidneys from IL-15 treated and non-treated mice at day 7 after adriamycin treatment. * shows podocyte feet effacement and glomerular basement membrane thickening. **(f)** Immunofluorescence of Podocalyxin (PODXL; red), a podocyte marker, and P62/SQSTM1 (green) in glomeruli from wild-type mice at day 7 after adriamycin injection with or without IL-15 treatment showing the accumulation of P62 in podocytes during Adriamycin model and decreased by IL-15 treatment. Nuclei were counterstained with Hoechst (blue). Figure subparts with prime indicate higher magnification. Bars=50 mm. **(g)** Associated quantification of the P62+ area expressed as the percentage of the P62/PODXL positive area. Values are presented as individual plots and mean ± SEM.

## Discussion

STAT5 proteins are classically described in immune cells. Here, we show that the STAT5B transcription factor is induced in podocytes in FSGS. We demonstrate that STAT5 has a protective effect in glomerular in 2 experimental models of glomerular disease. Moreover, exogenous IL-15 can activate STAT5, maintaining podocytic autophagy and conferring glomerular protection. Thus, IL-15 offers a potential new therapy for glomerular disease. The tissue-specific role of STAT5 was first recognized in 1994. This study demonstrated that STAT5 can be activated by prolactin in mammary glands^11^. Since then, several studies have further described the critical role of STAT5 proteins in adipogenesis, cellular differentiation, immune function and oncogenesis^12,25^. STAT5 also controls cell survival through the regulation of pro-survival genes such as *BCL-XL*^26^. Also, STAT5 is activated by numerous cytokines such as IL-15, erythropoietin, growth hormone, and prolactin. A protective role for STAT5 proteins has been shown in neuronal cells and in experimental hepatotoxicity, effects mediated through anti-apoptotic factors such as Bcl-xL^26^. With respect to the kidney, Fragiadaki *et al*. recently demonstrated that tubular STAT5 activation promotes cell proliferation and cyst formation in a model of polycystic kidney disease^27^ but no report was found related to STAT5 function in podocytes. Additionally, a recent study defined STAT5 as a key regulator of repair following experimental ischemic acute kidney injury^28^.

The finding that STAT5 is activated in human podocytes is novel. In glomerular disease, the degree of STAT5 activation appeared to be related to disease severity, as biopsies of patients with more severe collapsing variants of FSGS showed a greater STAT5 activation within enlarged nuclei. Interestingly, aberrant podocyte mitotic activity with enlarged nuclei has been described in FSGS^29–31^. STAT5 proteins could thus play a role in cellular proliferation, as suggested by Fragiadaki *et al* in tubular cells. Our preliminary results also show an upregulation of STAT5 in injured human tubular cells which also displayed nuclear enlargement which might be explained by tubular proliferation.

There are few pro-survival pathways described in podocytes. These include Peroxisome Proliferator Activated Receptor-Gamma (PPARγ), Kruppel like factors and BMP7.^3,7,32^ However, the vast majority of them do not have commercially available agonists and often exert opposing effects on glomerular and tubular cells leading to potential side effects if used in kidney diseases. For example, PPARγ can be activated by thiazolidinediones once used in diabetes.^33^ Unfortunately, thiazolidinediones are no longer commercially available due to their hepatotoxicity and cardiovascular side effects promoted by salt retention from the kidney. ^34,35^

Recent work on the STAT proteins in kidney disease have mainly focused on STAT3. It has been shown that STAT3 activation promotes deleterious effects both on podocyte (experimental glomerulonephritis and HIV-associated nephropathy [HIVAN])^7 8^ and on tubular cells by stimulating fibrosis.^9^ Compared to STAT3, STAT5 activation has an opposite (positive) effect in glomerular epithelium suggesting that a STAT3/STAT5 balance might exist in epithelial cells similar to that seen in immune cells. In lymphocytes, STAT3 promotes Th17 differentiation and represses Treg function whereas STAT5 promotes Treg differentiation and suppresses Th17 differentiation.^36^ STAT3 and STAT5 also have opposing effects on apoptosis in B lymphocytes.^37^ Therefore, we propose that a similar STAT3/STAT5 balance might exist in the kidney, and that this balance is upset in kidney disease. STAT5 activates or maintains an essential pro-survival pathway in the podocytes, that are terminally differentiated cells as autophagy.

We show here that IL-15 activates STAT5 in podocytes, as observed previously in immune cells, and alleviated glomerular injury maintaining autophagic flux. Autophagy activation by an exogenous compound in vivo is an unmet need that could be overcome by IL-15. IL-15, or its super agonist ALT-803, is commercially available with several ongoing clinical trials in cancer.^38,39^ Previously, we demonstrated that podocytes express IL-15 receptors during human crescentic glomerulonephritis.^1^ Further studies are needed to determine the minimal effective dose of IL-15 in a range of experimental models of kidney disease. Importantly, the kidney is a potential producer of IL-15 following injury, and this endogenous IL-15 can limit kidney damage. This is supported by data from global *Il15* knockout mice, which display accentuated tubular injury in experimental glomerulonephritis^19^. *In vivo*, a mix of IL-15/IL-15Rα has been shown to be more effective than IL-15 alone in several studies.^40^ By stabilizing IL-15, IL-15Rα fosters IL-15 transactivation through the IL2Rβ/γC receptor.^20^

We acknowledge some limitations of our study. First, the respective roles of STAT5A and STAT5B in the kidney epithelium are not fully understood. Although we observed strong activation of STAT5B in human kidney, additional studies are needed to resolve this point. Our *in vivo* studies cannot differentiate between the respective roles of STAT5A and STAT5B as the loci for both were deleted in the original construct designed by Hennighausen et al^15^.

Altogether, our data demonstrate that STAT5 is activated in podocytes during FSGS and has an important protective role activating autophagy. Activating podocytic STAT5 represents a new therapeutic avenue with the potential for a range of beneficial effects in glomerular disease.

## Author contributions

A NIASSE: Conceptualization, Methodology, Validation, Investigation, Formal analysis, Writing, Visualization; K LOUIS: Methodology, Investigation; O. LENOIR: : Investigation, Methodology, Writing; C SCHWARZ: Methodology, Investigation, Writing; X XU: Methodology, Investigation; A COUTURIER: Methodology, Investigation; H DOBOSZIEWICZ: Methodology, Investigation; A CORCHIA: Methodology, Investigation; S PLACIER: Investigation, Methodology; S VANDERMEERSCH: Investigation, Methodology; L HENNIGHAUSEN: Resources; P FRERE: Software, Investigation, Formal analysis; P GALICHON: Investigation, Methodology; B SURIN: Investigation; S OUCHELOUCHE: Investigation; L LOUEDEC: Resources, Investigation; T MIGEON: Investigation; MC VERPONT: Investigation, Visualization; D BUOB: Resources, Methodology; YC XU-DUBOIS: Investigation, Visualization; H FRANCOIS: Conceptualization, Methodology; E RONDEAU: Conceptualization, Supervision, Resources, Writing; L MESNARD: Conceptualization, Methodology, Supervision, Writing; J HADCHOUEL: Investigation, Methodology, Design, Supervision, Writing; Y LUQUE: Investigation, Conceptualization, Methodology, Supervision, Writing, Project administration, Funding acquisition

## Data and materials availability

The authors declare that the data supporting the findings of this study are available within the paper and its supplementary information files. All data are available from the corresponding authors upon reasonable request.

## Supplementary Materials

**Supplementary Methods**

**Supplemental Figure 1**. STAT5 deficiency in podocytes does not affect renal development in mice

**Supplemental Figure 2**. STAT5 deficiency in podocytes has no influence on depletion of blood cells induced by adriamycin treatment

**Supplemental Figure 3**. Interleukin-15 immune effect in adriamycin-induced nephropathy

**Supplemental Table 1**. Primers used for RT-qPCR

**Supplemental Table 2**. Patient’s characteristics

## Acknowledgements

This work was supported by Institut National de Santé et de la Recherche Médicale (Inserm), Sorbonne Université and a research grant from the French Agence Nationale de la Recherche (PodoGammaC grant ANR JC to YL). We are grateful to Assistance Publique - Hôpitaux de Paris (Médaille d’argent et Fonds d’Etudes et de Recherche du Corps Médical) and the Société Francophone de Néphrologie, Dialyse et Transplantation (SFNDT) for supporting Y.L. We thank Liliane Louedec, Jessy Renciot, Claude Kitou, and Gaetan Girault for assistance in animal care and handling, Nicolas Sorhaindo (CRI – Plateforme de Biochimie UMR 1149 Université Paris Diderot, Paris, France) and Gaelle Brideau (Renal Metabolism and Physiology Laboratory, Centre de Recherche des Cordeliers, UMRS 1138, ERL 8228, Paris, France) for biochemical measurements. We acknowledge excellent administrative support from Mélanie Charlery and Emmanuelle Bénard. XX, HD, CS, were funded by Société Francophone de Néphrologie Dialyse et Transplantation (SFNDT). This work has benefited from the facilities and expertise of the CoRaKiD histology and imaging core facilities (Common and Rare Kidney Diseases Unit, INSERM UMR_S1155, Paris, France).

