## Supplementary material for "Protective Role of Podocytic IL-15 / STAT5 Pathway in Experimental Focal and Segmental Glomerulosclerosis": Supp Materials BioRxiv.pdf

#### Supplementary Materials

##### Supplemental Figure 1. STAT5 deficiency in podocytes does not affect renal development in mice

**(a)** Schematic representation of the generation of mice with podocyte STAT5 deficiency obtained by mating mice expressing the Cre recombinase under the podocin promoter (*Nphs2*.Cre mice) with mice expressing *Stat5* alleles with loxP sites flanking exon (*STAT5*$^{lox/lox}$ mice). **(b)** Representative images of Cre recombinase staining (upper panel, scale bar: 50 µM), Masson's trichrome (middle panel, scale bar 100µM) staining and transmission electron microscopy (lower panel, scale bar: 2 µM) in kidneys of 10-weeks-old *Stat5*$^{\text{lox/lox}}$ and *Nphs2*.Cre-*Stat5*$^{\text{lox/lox}}$ mice. **(c)** BUN and urine albumin-to-creatinine ratio in 10-week-old *Stat5*$^{\text{lox/lox}}$ and *Nphs2*.Cre-*Stat5*$^{\text{lox/lox}}$ mice. Individual values are shown as dots. Bars represent means.

##### Supplemental Figure 2. STAT5 deficiency in podocytes has no influence on depletion of blood cells induced by adriamycin treatment

(**a**) Body weight evolution in *Stat5*$^{\text{lox/lox}}$ *and Nphs2*.cre *Stat5*$^{\text{lox/lox}}$ mice after adriamycin or vehicle treatment. n=4 to 10 mice per group. Data represent means. **(b)** Hemoglobin level, total leukocyte count, platelet count, lymphocyte count and neutrophil count in the blood of *Stat5*$^{\text{lox/lox}}$ *and Nphs2*.cre-*Stat5*$^{\text{lox/lox}}$ mice at day 7 after vehicle (sham) or adriamycin treatment Individual values are shown in dots. Bars represents means. ****$p < 0.0001$. **p<0.01. *p<0.05.

##### Supplemental Figure 3. Interleukin-15 immune effect in adriamycin-induced nephropathy

**(a)** Representative images of the spleen of WT or IL-15 treated mice after adriamycin treatment. **(b)** Representative images of CD3 (upper panel) and F4/80 (lower panel) staining in kidneys from IL-15 treated and non-treated mice at day 7 after adriamycin or vehicle treatment (Scale bar : 2µm). **(c)** Quantification of F4/80 staining in kidneys (left panel)**.** Individual values are shown and the bars correspond to the means. 5 fields per mice were analyzed with Image J. RT-qPCR quantification of the expression of *Mcp1* (right panel) in the kidneys from 12-week-old BALB/C mice at day 7 after adriamycin injection with or without IL-15 treatment. **(d)** Quantification of CD3 staining in kidneys (left panel)**.** Individual values are shown and the bars correspond to the means. 5 fields per mice were analyzed with Image J. **p<0.01 and *p<0.05.

**Supplemental Table 1. Primers used for RT-qPCR**

**Supplemental Table 2. Patients characteristics**

**Supplementary methods**

*Western blot analysis*

Proteins were extracted from primary cultured mouse podocytes, mouse whole kidney or cultured human podocytes with RIPA buffer with protease and phosphatase inhibitor cocktail as described above. Total protein concentration was measured using the Bradford assay (Biorad). Samples were fractionated by SDS-PAGE using NuPAGE 12% gels (Invitrogen) under reducing conditions and then transferred to a nitrocellulose membrane. Membranes were incubated in TBST (TBS 1X with 0.1% Tween) with the appropriate primary antibodies: anti-phosphorylated STAT5 (Cell Signaling, D47E7, 1:1000), anti-phosphorylated STAT5B (Abcam, ab52211, 1:1000), anti-STAT5 (Abcam, ab16276, 1:1000). GAPDH (Sigma-Aldrich, 1:40000), rabbit anti-LC3 (1:1000, #3868, Cell Signaling). Anti-tubulin (1:5000, Abcam, ab6160) or anti-beta-actin (NB100-56874, Novusbio, 1:1000) antibodies were used as loading control. ImageJ software (National Institutes of Health) was used for quantification.

*Cell Culture*

The laboratory of Moin Saleem initially generated AB8/13 human podocytes[1]. They were cultured in Roswell Park Memorial Institute-1640 (RPMI-1640)-based medium supplemented with 10% fetal bovine serum (FBS; Biosera),1% Insulin-Transferrin-Selenium (Invitrogen, Breda). Podocytes were cultured at 33°C in a 5% $CO_2$ incubator and differentiated at 37°C for 10-14 days. Recombinant human IL-15 (R&D systems) and recombinant murine IL-15Rα-Fc (R&D systems), both resuspended in phosphate buffered saline (PBS), were mixed and incubated for 30 min at 37°C. Following serum starvation, podocytes were cultured in the presence or the absence of IL-15/IL-Rα with or without bafilomycin A1 (100 nmol/L, Enzo Life Sciences, BML-CM110-0100) for 4 hours.

*Generation of STAT5B knock-out in cultured human podocytes*

CRISPR/Cas9-mediated *STAT5B* gene knockout was performed in AB8/13 human podocytes. A total of $4 \times 10^5$ cells (plate 6 wells) were transfected with the Cas9 gene, puromycin reporter gene, and single guide RNA (sgRNA) targeting *STAT5B* gene exon 2 (ATCAGATGCAAGCGTTATA) expression vectors (PX459 *hSTAT5B*) using the PEI reagent (1mg/ml) according to the manufacturer's instructions. Two days after transfection, puromycin selection was performed in plate 6 wells by culturing the transfected cells in the presence of 5 mg/ml of puromycin (5 µg/ml final) for two days. Single cell clones from the puromycin-selected cells were generated by limiting dilution into 96-well plates at a density of 0.5 cells/well in the absence of puromycin. The *STAT5B*-knockout clones with a biallelic mutation on the endogenous *STAT5B* gene were then selected by DNA sequencing and confirmed by Western blot analysis. Human podocytes and GS-knockout clones were maintained in T-25 flasks in a 5% $CO_2$/air mixture, humidified at 37°C. The medium for culture maintenance was Dulbecco's modified Eagle's medium (DMEM, Gibco, Grand Island, NY) supplemented with 10% fetal bovine serum (FBS, Biosera).

*IL-15 therapy protocol in vivo*

Recombinant human IL-15 (R&D systems) and recombinant murine IL-15Rα-Fc (R&D systems), both resuspended in phosphate buffered saline (PBS), were mixed and incubated for 30 min at 37°C. Each BALB/C mouse of the IL-15/IL-15Rα group and IL-15/IL-15Rα + adriamycin was treated with 2.5µg of IL-15 pre-complexed with 7.5 µg of the recombinant mouse IL-15Rα-Fc chimeric molecule (R&D Systems Europe Ltd, Abingdon, Oxon, UK) in 200 µl of PBS. Control mice were treated with 200 µl of PBS. All treatments were administered subcutaneously 1 day before adriamycin injection and then every two days until euthanasia.

### SUPPLEMENTAL FIGURE 1

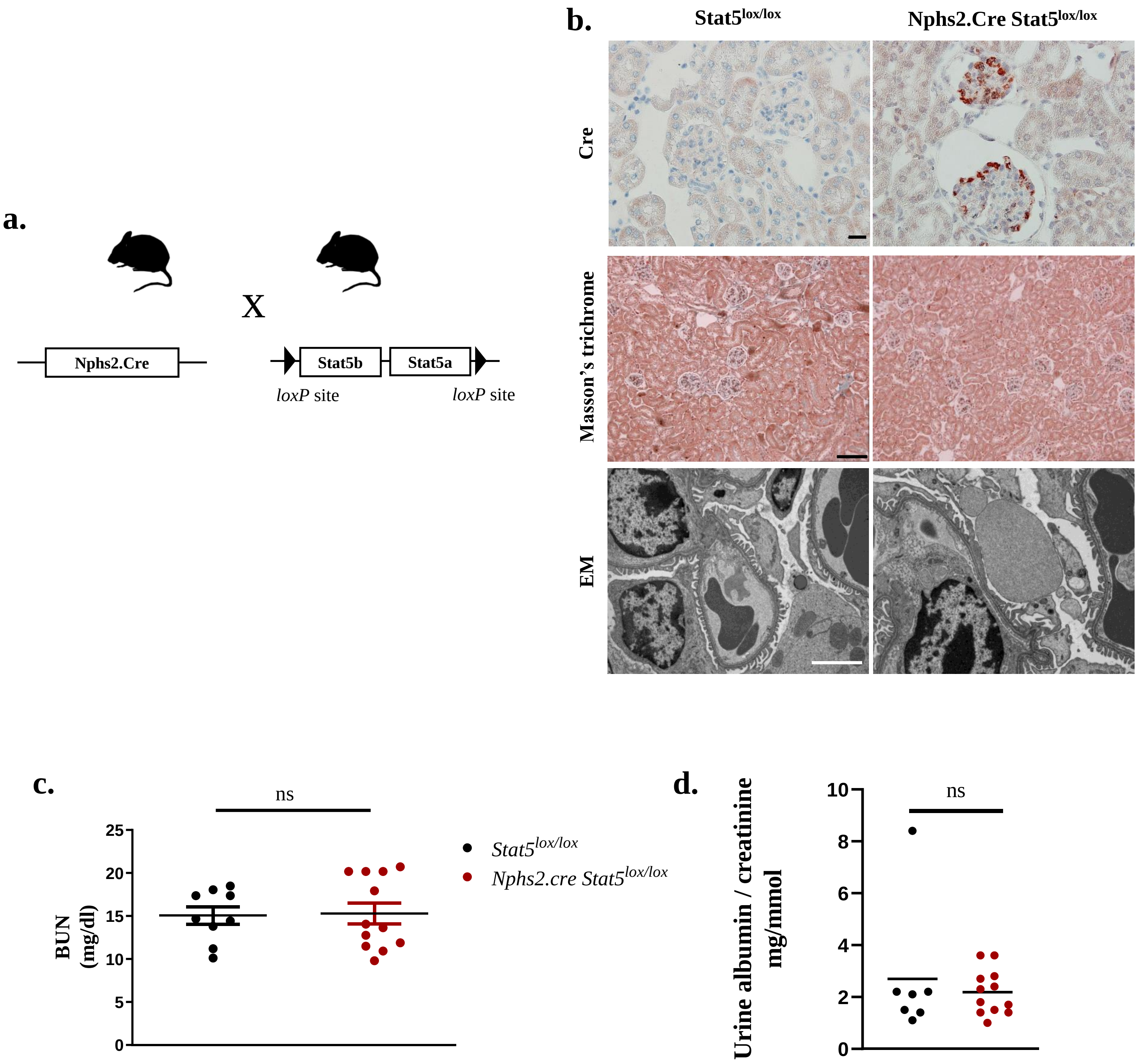

### SUPPLEMENTAL FIGURE 2

**a.**

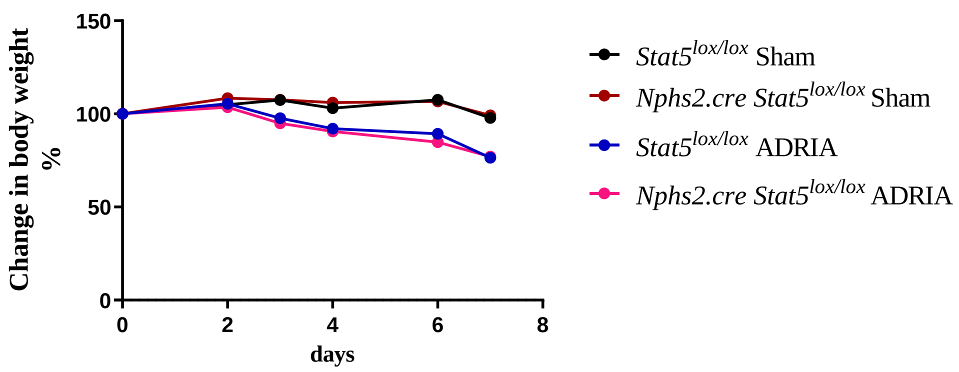


**b.**

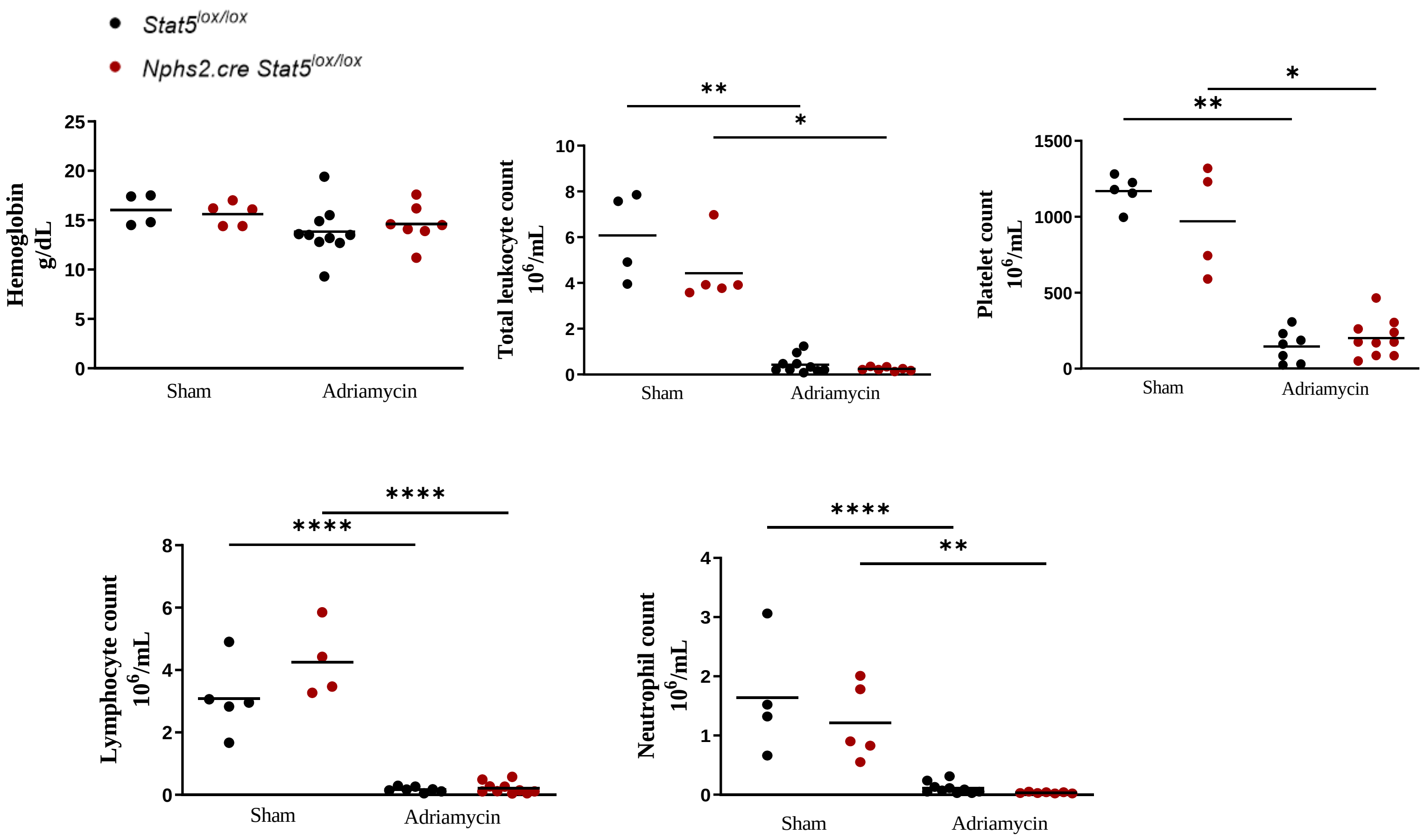

### SUPPLEMENTAL FIGURE 3

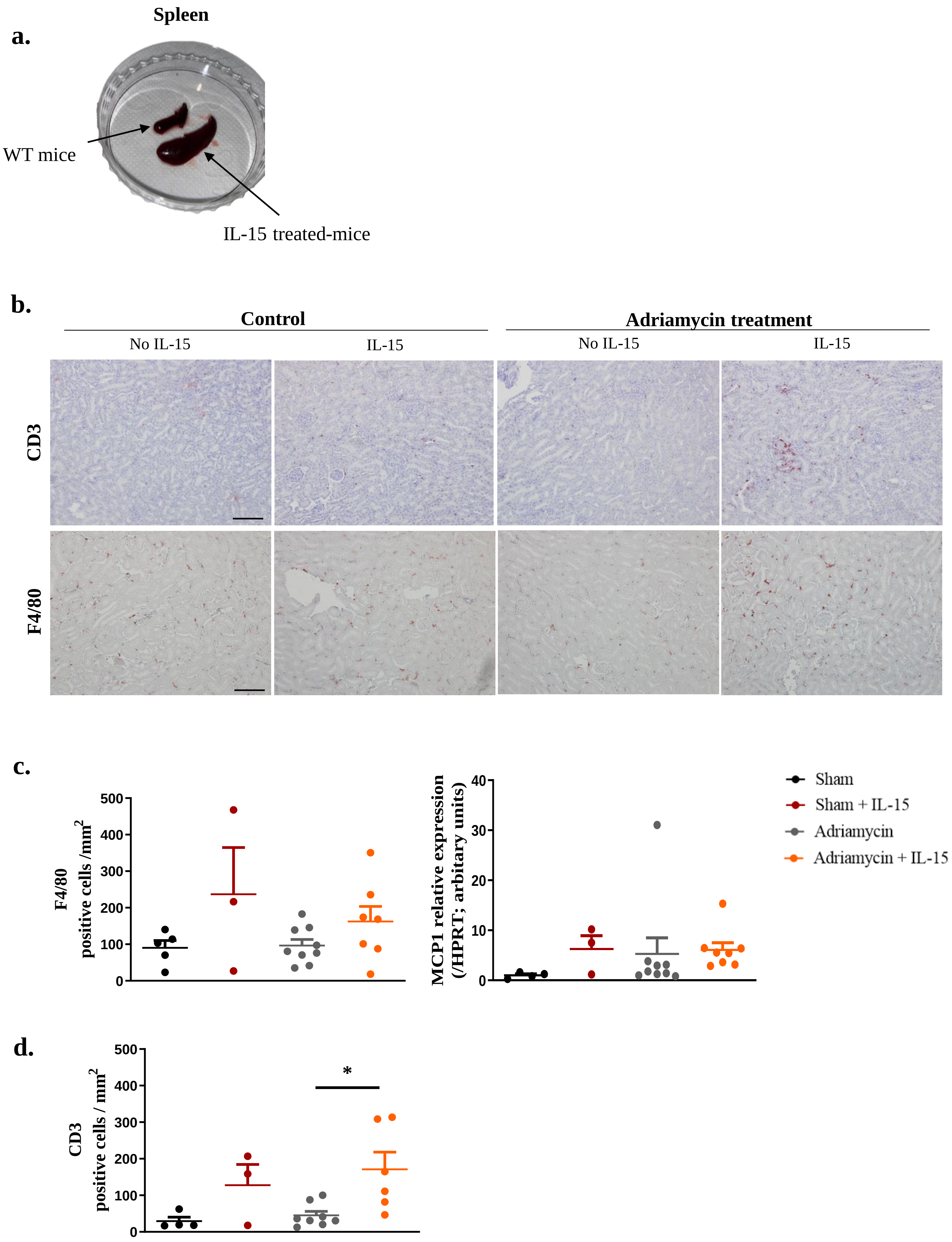

**Supplemental Table 1 : Primers used for RT-qPCR**

| **Target Gene** | **Forward primer** | **Reverse primer** |
|---|---|---|
| ***Mcp1*** | GGCTGGAGAGCTACAAGAGG | CTCTTGAGCTTGGTGACAAAAA |
| ***Hprt*** | GCAGCGGTAGCACCT | CTGGTTCATCATCGCTAATCA |
| ***Stat5a*** | TCTACGTGTTCCCAGACCGA | GCATTGACGAACCAAGTACAGG |
| ***Stat5b*** | CTTGTACGGCCAGCATTTCC | CAAGATCTATTGAGTCCCAGGC |

**Supplemental Table 2 : Patient's characteristics**

| Patient | Diagnosis | Age at kidney biopsy (years) | Sex | Plasma albumine (g/L) | Urine protein to creatinine ratio (g/mmol) | Plasma creatinine (µmol/L) | Other |
|---|---|---|---|---|---|---|---|
| 1 | HIVAN | NA | NA | NA | NA | NA | |
| 2 | HIVAN | 64 | M | 18 | 0,75 | 970 | |
| 3 | HIVAN | 31 | F | 18 | 2 | 922 | |
| 4 | Collapsing FSGS | 66 | M | 22 | 0,37 | 1070 | Malaria |
| 5 | Collapsing FSGS | 45 | M | 12 | 2,29 | 981 | Malaria |
| 6 | Collapsing FSGS | 38 | M | 20 | 1,12 | 676 | Parvovirus B19 |
| 7 | Collapsing FSGS | 53 | M | 33 | 1,09 | 738 | HHV-6 |
| 8 | FSGS | 34 | M | 36 | 0,21 | 76 | Obesity |
| 9 | FSGS | 29 | F | 32 | 0,3 | 83 | NA |
| 10 | FSGS | 37 | M | 43 | 0,06 | 126 | HT-high risk *APOL1* |
| 11 | Control | 27 | F | Normal | < 0,02 | 103 | Kidney Tx 3m |
| 12 | Control | 63 | M | Normal | 0,01 | 177 | Kidney Tx 3m |
| 13 | Control | 45 | F | Normal | 0,03 | 164 | Kidney Tx 3m |
